# Quantifying forms and functions of intestinal bile acid pools in mice

**DOI:** 10.1101/2024.02.16.580658

**Authors:** Koichi Sudo, Amber Delmas-Eliason, Shannon Soucy, Kaitlyn E. Barrack, Jiabao Liu, Akshaya Balasubramanian, Chengyi Jenny Shu, Michael J. James, Courtney L. Hegner, Henry D. Dionne, Alex Rodriguez-Palacios, Henry M. Krause, George A. O’Toole, Saul J. Karpen, Paul A. Dawson, Daniel Schultz, Mark S. Sundrud

**Author notes:** **Correspondence:** Mark S. Sundrud, Ph.D., Dartmouth Health and Geisel School of Medicine at Dartmouth, 1 Medical Center Drive, Lebanon, NH, 03756, USA.

## Abstract

Bile acids (BAs) are gastrointestinal metabolites that serve dual functions in lipid absorption and cell signaling. BAs circulate actively between the liver and distal small intestine (*i.e*., ileum), yet the dynamics through which complex BA pools are absorbed in the ileum and interact with intestinal cells *in vivo* remain ill-defined. Through multi-site sampling of nearly 100 BA species in individual wild type mice, as well as mice lacking the ileal BA transporter, Asbt/*Slc10a2*, we calculate the ileal BA pool in fasting C57BL/6J mice to be ∼0.3 μmoles/g. Asbt-mediated transport accounts for ∼80% of this pool and amplifies size, whereas passive absorption explains the remaining ∼20%, and generates diversity. Accordingly, ileal BA pools in mice lacking Asbt are ∼5-fold smaller than in wild type controls, enriched in secondary BA species normally found in the colon, and elicit unique transcriptional responses in cultured ileal explants. This work quantitatively defines ileal BA pools in mice and reveals how BA dysmetabolism can impinge on intestinal physiology.

## Introduction

The mammalian gastrointestinal (GI) tract is now known to harbor more than 100 bile acid (BA) species, each possessing unique physicochemical properties^1^. As such, increasingly advanced analytical methods (*e.g*., metabolomics) are being developed to quantify growing numbers of BA species within complex biospecimens (*e.g*., blood, stool) ^2,3^. Whereas these studies have transformed our understanding of BA biochemistry, even the most powerful instruments used for single-site measurements fail to inform broader regulatory dynamics that underlie BA circulation and signaling *in vivo*.

Hepatocytes synthesize a quantitatively large, but compositionally simple, pool of primary BAs [a.k.a., bile salts; cholic acid (CA), chenodeoxycholic acid (CDCA) in humans; CA, CDCA, α-muricholic acid (αMCA), and β-muricholic acid (βMCA) in mice] via cholesterol catabolism^4^. These BAs are conjugated to amino acids [*e.g*., glyco (g)CA, tauro (t)CA], transported into bile ducts, concentrated in the gallbladder, and secreted into the duodenum. BA n-acyl amidation limits membrane permeability and cytotoxicity and restricts BAs to the lumen of the proximal small intestine (SI) for efficient micelle formation^4,5^.

As most BA-dependent lipid and fat-soluble vitamin (*e.g*., Vit. A/D/E/K) absorption occurs in the proximal- and mid-small intestine (SI; *i.e*., duodenum and jejunum, respectively), an increasing proportion of BAs reaching the distal SI (*i.e*., ileum) are found in free/non-micellular forms. It is estimated that ∼95% of free BAs are reclaimed from the ileal lumen by specialized enterocytes expressing the apical sodium-dependent bile acid transporter (ASBT/*SLC10A2*) ^4,6^. Intra-epithelial BAs are bound by the ileal BA-binding protein (iBABP; *e.g*., FABP6), trafficked to the basolateral surface, and transported into the underlying mucosal tissue (*i.e*., lamina propria) by the OSTα/β complex^7,8^. Because of active absorption, immune and epithelial cells in the ileum are exposed to high, potentially toxic BA concentrations^9,10^. Ileal enterocytes manage BA toxicity via the nuclear receptor (NR), FXR/*Nr1h4*, which upon activation by intracellular BAs promotes expression of several adaptive genes, including *Fapb6* (to increase BA-binding capacity), OSTα/β (*SLC51A/SLC51B*; to increase basolateral membrane efflux capacity), and *Fgf15* (*FGF19* in humans; to restrict hepatic BA synthesis) ^11-18^. In parallel, immune cells in the ileal lamina propria leverage an orthogonal NR, CAR/*Nr1i3*, to activate a ‘hepatocyte-like’ transcriptional response upon BA exposure that limits oxidative stress and inflammation^19,20^. Ultimately, ileal BAs enter portal capillaries to be carried back to the liver for reuptake, potential enzymatic modification and re-secretion into bile, thus beginning another enterohepatic cycle^4,21^.

The fraction of primary BAs escaping ileal absorption (∼5% at homeostasis), enter the large intestine (LI; *i.e*., cecum and colon) and are increasingly subject to metabolism by commensal microbiota^1^. The ‘gatekeeper’ reaction in bacterial BA metabolism is deconjugation [*i.e*., deamidation; (t)CA→CA or (t)CDCA→CDCA] by bile salt hydrolase (*bsh*) enzymes^22^. Following deconjugation, primary BAs can undergo diverse microbe-dependent reactions including dehydroxylation, epimerization of hydroxyl groups to form “iso” bile acids, oxidation and reduction to form “allo” bile acids, secondary side chain conjugation to other amino acids (*e.g*., leucine, tyrosine, phenylalanine), and esterification at the 3-hydroxy position to fatty acids^1,23-25^. Removal of C-7 hydroxyl groups (*i.e*., 7α-dehydroxylation) is among the most common and important microbial BA biotransformation, producing major secondary BAs deoxycholic acid (DCA, from CA) and lithocholic acid (LCA, from CDCA), and utilizing enzymes encoded by the bile acid-induced (*bai*) operon, a common genetic trait of several commensal genera (*e.g*., *Clostridia, Bacillus*) ^26^. The LI BA pool is thus quantitatively smaller, but also more diverse and hydrophobic, than the SI BA pool. Colonic BAs are either excreted in feces or passively absorbed for portal recirculation to the liver. BAs diffusing across the colonic epithelium interface with mucosal immune cells, where LCA and several of its metabolites have been shown to interact with still other NRs – VDR/*Nr1i1*, RORγt/*Nr1f3* – and promote immune tolerance^27,28^.

In line with their diverse and critical functions in GI health, changes in the circulation and/or metabolism of BAs have been linked to a growing list of human diseases, from metabolic dysfunction-associated steatotic liver disease (MASLD) and GI cancers, to inflammatory bowel disease (IBD) and cystic fibrosis (CF) ^29,30^. However, deciphering the underlying causes and functional consequences of disease-associated BA dysmetabolism has proven more formidable. In IBD and CF, for example, levels of secondary BAs in stool are often found to be reduced in patients *vs*. controls^31-34^. While these results are generally interpreted as indicative of decreased bacterial BA metabolism due to dysbiosis^1,35^, alternative explanations cannot be excluded without augmented information about hepatic synthesis and/or intestinal absorption. Functionally, it is presumed that exposure of host intestinal cells to disease-associated BA pools can precipitate mucosal immune and epithelial dysfunction, yet it remains uncertain how BA concentrations in clinically-accessible sites (*e.g*., stool, serum), or even in the intestinal lumen, correspond to local levels in intestinal tissues – the latter being limited by variable and undefined rates of active and passive absorption *in vivo*. Finally, even if tissue-associated BA concentrations can be extrapolated or measured empirically, it is not known how complex pools of BAs compete for interactions with host cells and receptors.

Appreciating that BA homeostasis is an active process, governed by higher-order endocrine connections between the liver (where BAs are made), ileum (where most BAs are absorbed) and colonic microbiota (where most BAs are metabolized), we sought to develop an integrated methods set to quantify, and functionally interrogate, ileal BA absorption in mice. We describe a ‘multi-GI-omics’ platform that features parallel sampling of nearly 100 BA species in three GI sites of individual mice, and that yields quantitative insights into ileal BA absorption. Combining metabolomics with metagenomics informs aspects of BA dysmetabolism that involve altered microbial function. Incorporating host gene expression analyses – particularly in the liver, ileum and colon – reveals whether, where and how BA-dependent signaling is affected *in vivo*. Finally, molecular functions of intestinal BA pools can be examined independent of confounding *in vivo* variables (*e.g*., on microbiota, host metabolism) using cultured intestinal explants. We validate these approaches using mice that lack the ileal BA reuptake transporter, Asbt, and that harbor a profoundly different enterohepatic BA pool. In the process, we provide working answers to long-standing questions about the absorption and functions of the ileal BA pool. This work reshapes extant views of the ‘gut-liver axis’, moves BA understanding away from individual species and towards physiological pools, and forges new paradigms for deciphering the forms and functions of endogenous BAs in human health and disease.

## Results

### A system-level perspective of *in vivo* BA circulation and metabolism

To leverage the strengths of mass spectrometry, while also appreciating its limitations (*e.g*., lack of spatial resolution in intestinal tissues), we surveyed nearly 100 BA species and metabolites (representative chromatogram shown in **Figure S1**) in three parallel GI sites of individual mice, including: (*i*) SI luminal content (siLC) – to define the pool of BAs made by the liver and secreted into the intestine; (*ii*) superior mesenteric vein blood (smvB) – to resolve the pool of SI luminal BAs reabsorbed primarily in the ileum; and (*iii*) feces – to capture the fraction of BAs that escape ileal absorption and become metabolized in the LI (**Figure 1A**). We used fecal BAs as a surrogate for those in the colon lumen because pilot studies revealed these two pools were highly similar (**Figure S2A**). A fourth site, peripheral blood (pB), was included to formally establish how BAs in peripheral circulation – an increasingly common clinical measurement – reflect intestinal pools. To avoid inter-animal variation in post-prandial BA secretion and ileal delivery^36^, mice were fasted prior to sample collection.

**Figure 1.**
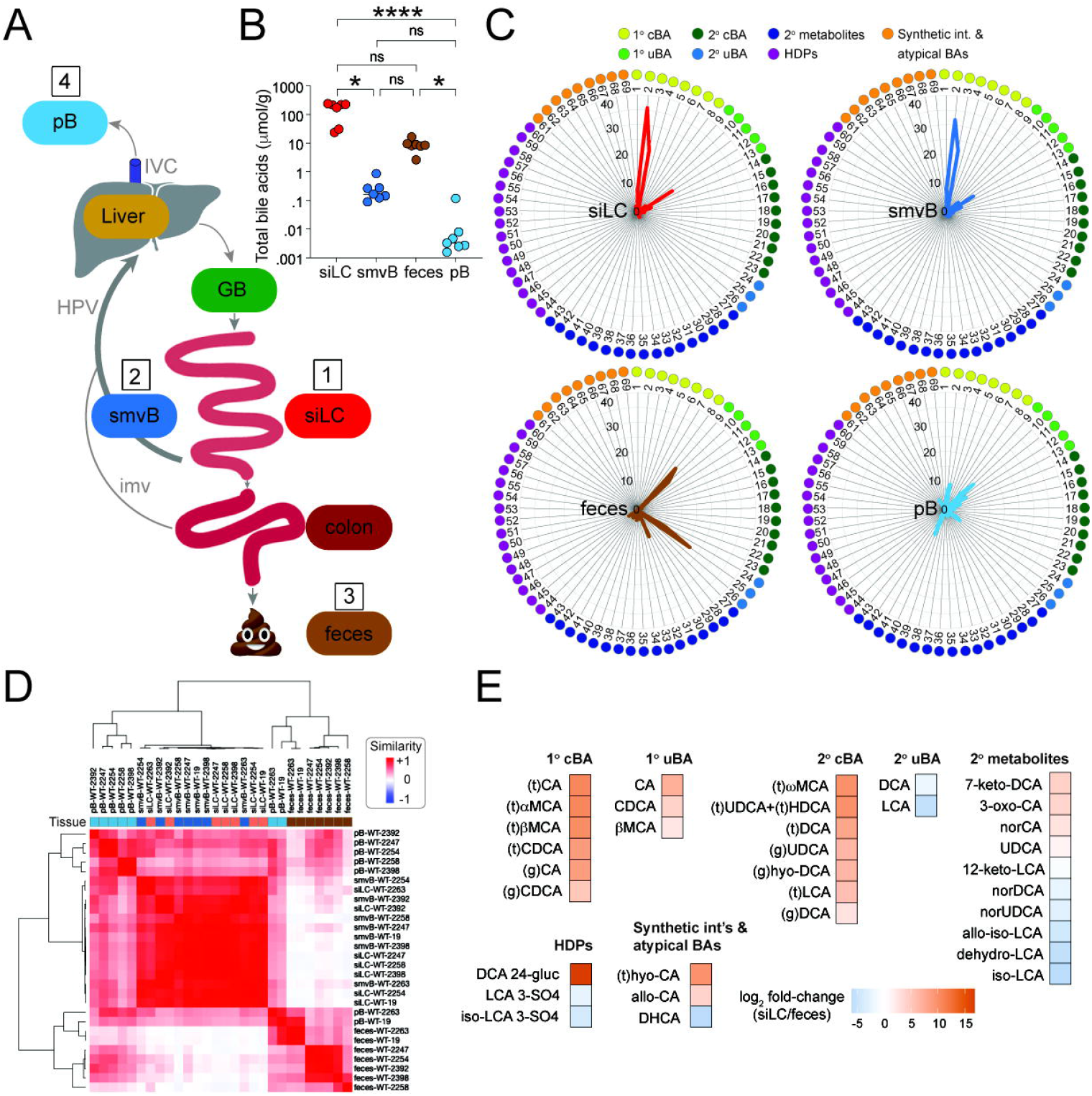
Multi-site analysis of *in vivo* bile acid metabolomes. **(A)** A schematic diagram of bile acid (BA) enterohepatic circulation adapted from^59^. GB, gallbladder; HPV, hepatic portal vein; imv, inferior mesenteric vein; IVC, inferior vena cava; pB, peripheral blood; siLC, small intestine luminal contents; smvB, superior mesenteric vein blood. 88 BA species and metabolites were analyzed by LC-MS/MS in the sites numbered 1-4. **(B)** Total BA concentrations (μmol/g of luminal content or plasma; *n* = 7), determined by LC-MS/MS, in the 4 sites numbered and color-coded in (A) from C57BL/6J wild type (B6) mice. Data are presented as the sum of 69 reliably detectable species described in Figure S2B-C. Each symbol represents data from one mouse; horizontal lines indicate means. \**P* < 0.05, \*\*\*\**P* < 0.0001, Friedman test with Dunn correction for multiple comparisons ; ns, not significant. **(C)** Percentages of each BA species within the total pool from the 4 sites numbered and color-coded in (A) of wild type B6 mice. The 69 reliably detectable species presented here are listed, grouped and color-coded as in Figure S2C. Mean values are from 7 mice. 1° cBAs, primary conjugated BAs; 1° uBAs, primary unconjugated BAs; 2° cBAs, secondary conjugated BAs; 2° uBAs, secondary unconjugated BAs; 2° metabolites, secondary metabolites; HDPs, hepatic phase 2 detoxification products. **(D)** Similarity matrix of endogenous BA pools in the 4 sites numbered in (A) from wild type B6 mice. Calculated based on levels of the 69 reliably detected BA species described in Figure S2B-C. Sample IDs list tissue type, genotype and cage number; tissues are color-coded at top. **(E)** Heat maps of the relative abundance (Log_2_ fold-change) of individual BA species, determined as above by LC-MS/MS, that are significantly different (*P*adj < 0.05, Wilcoxon rank-sum test) between siLC and feces.

Liquid chromatography coupled with tandem mass spectrometry (LC-MS/MS) quantified 88 discrete BA species and metabolites in the above-four sites of 7 specific pathogen free (SPF) C57BL/6J (B6) wild type mice house in different cages. On analysis, 69 of 88 species displayed median values >0 in at least one of the three GI sites and were considered ‘reliably detectable’ (**Figure S2B**). These BA species and metabolites became the focus of downstream analyses and clustered into 7 general classes based on likely biosynthetic origins (**Figure S2C**) ^26,37,38^: (*i*) primary conjugated BAs (1° cBAs), which constitute most BAs synthesized and secreted by the liver at homeostasis; (*ii*) primary unconjugated BAs (1° uBAs), which reflect products of bacterial 1° cBA deconjugation prior to further microbial or host metabolism; (*iii*) secondary conjugated BAs (2° cBAs), which are secondary unconjugated BAs re-conjugated in hepatocytes following uptake from the intestines; (*iv*) secondary unconjugated BAs (2° uBAs), which are canonical 7α-dehydroxylation products generated from 1° uBAs via LI bacterial metabolism; (*v*) secondary (2°) metabolites, which are more extensively modified 2° uBAs (*e.g*., iso-LCA, 3-oxo-CA, etc.) produced via bacterial metabolism; (*vi*) hepatic phase 2 detoxification products (HDPs), including sulfate and glucuronide conjugates that are produced primarily in hepatocytes to limit BA toxicity and promote clearance; and (*vii*) synthetic intermediates and atypical BAs, which are normally secreted in trace amounts by healthy livers, but can increase during active liver disease (*e.g*., hepatobiliary disease, cancer) ^39-42^.

On aggregate, BAs were enriched in enterohepatic tissues (*i.e*., siLC, smvB) – as expected – reaching fasting levels of ∼160 μmol/g in siLC and ∼0.16 μmol/g (∼200 μM) in smvB (**Figure 1B**). siLC and smvB BA pools were compositionally akin to each other, consistent with efficient ileocecal absorption in the mouse, whereas both pools were clearly distinct from those in matched fecal samples (**Figure 1C-D**). 1° cBAs and 1° uBAs, as well as 2° cBAs, were enriched in enterohepatic tissues, whereas 2° uBAs (*e.g*., DCA, LCA) and 2° metabolites (*e.g*., iso-LCA, alloiso-LCA) were augmented in feces (**Figure 1E**). Importantly, pB BAs were the most variable between animals, displayed the highest α-diversity (*i.e*., intra-sample complexity), and presented mixed features of enterohepatic and fecal pools (**Figure 1C-D, S2D**). This result is consistent with the anatomical source, as metabolites in pB include those that evade hepatic first-pass clearance following portal recirculation from both the SI (via the SMV) and LI [via the inferior mesenteric vein (IMV)] (**Figure 1A**). Correlation matrices generated for all 69 BAs in each of the four sites revealed that fecal BAs had the fewest quantitative relationships with BAs in other sites (**Figure S2E**), further highlighting *in vivo* compartmentalization of SI (enterohepatic cycling) and LI BA pools.

### ASBT-deficiency reveals interconnected response networks underlying BA homeostasis

To explore the validity and utility of this approach further, we introduced a genetic perturbation – Asbt ablation – that drives BA dysmetabolism in robust and predictable ways^43^. The same analyses performed on wild type or Asbt-deficient (*Slc10a2*^-/-^) mice reared together until the day of analysis reproduced previous findings that loss of Asbt transport results in ∼5-fold fewer BAs in enterohepatic tissues (siLC, smvB), as well as commensurate ∼5-fold increases in fecal BA excretion (**Figure 2A**). Perhaps more interestingly, the remnant pool of enterohepatically circulating BAs in Asbt-deficient mice displayed increased α-diversity, compared with wild type controls, and was compositionally indistinguishable from both wild type and Asbt-deficient fecal pools (**Figure 2A-C**). At the species level, reshaping of the enterohepatic BA pool in Asbt-deficient mice was due to the depletion of virtually all 1° cBAs and 1° uBAs – consistent with previous ideas that 1° cBAs are preferred Asbt transport substrates in the ileum^44^ – and concomitant increases in 2° uBAs and metabolites normally enriched in the LI (**Figure 2D**). These results suggest that Asbt-mediated ileal BA absorption sustains both the unique size and composition of the enterohepatic BA pool.

**Figure 2.**
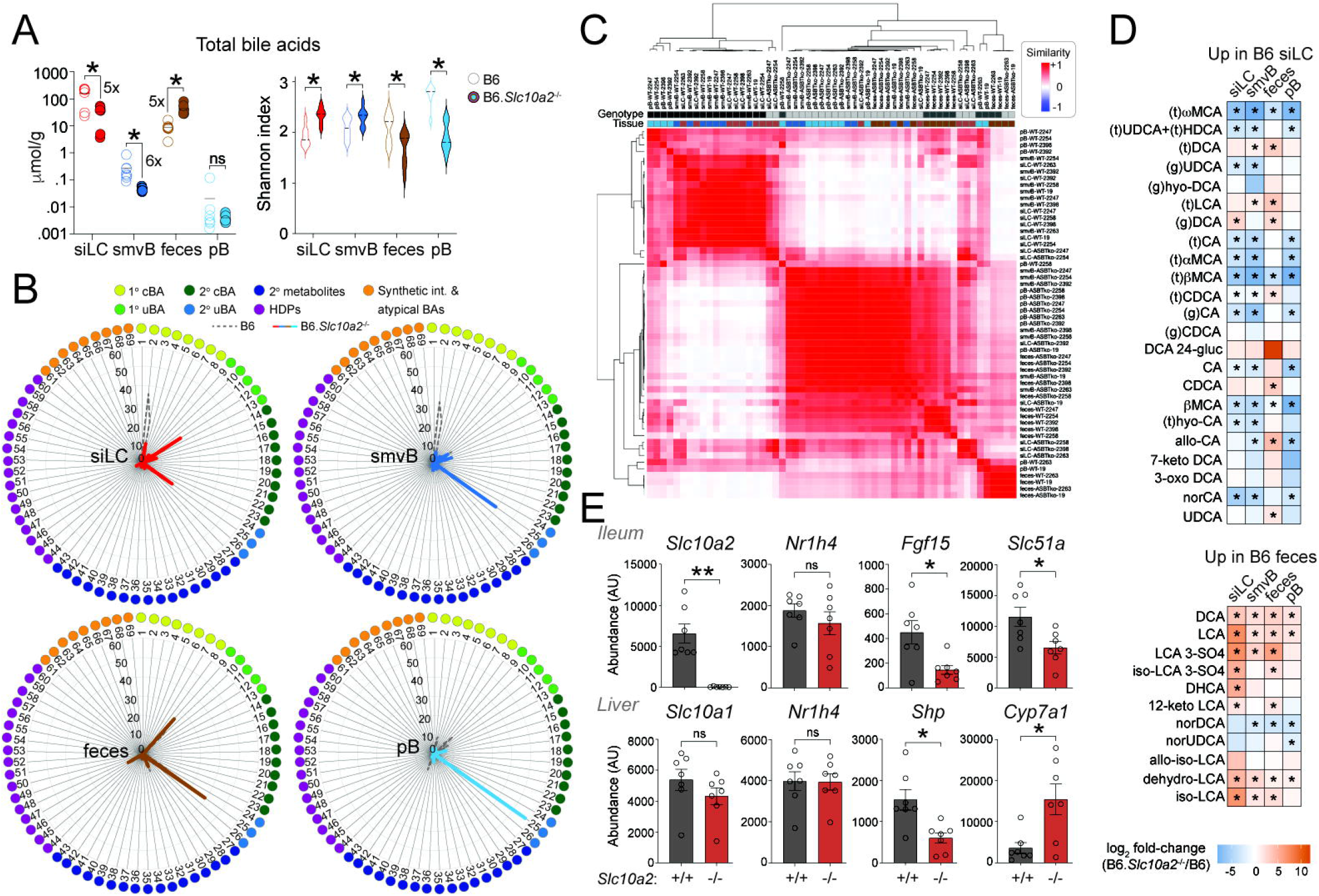
Asbt is required for *in vivo* compartmentalization of enterohepatic and fecal bile acid pools. **(A)** *Left*, total bile acid (BA) concentrations (μmol/g of luminal content or plasma; *n* = 7), determined by LC-MS/MS, in the 4 sites numbered and color-coded in Figure 1A from C57BL/6J wild type (B6, open circles) or Asbt-deficient (B6.*Slc10a2*^-/-^, filled circles) mice. Data shown are sums of 69 reliably detectable BA species described in Figure S2B-C. Each symbol represents data from one mouse; horizontal lines indicate means. *Right*, violin plots of α-diversity (Shannon index; *n* = 7) within indicated BA pools of wild type or Asbt-deficient mice analyzed as above. Data from wild type are the same as in Figure 1B (for total BA concentrations) and Figure S2D (for α-diversity). pB, peripheral blood; siLC, small intestine luminal contents; smvB, superior mesenteric vein blood. \**P* < 0.05, paired two-tailed Student’s *t* test; ns, not significant. **(B)** Percentages of each BA species within the total pool from the indicated sites of wild type (B6, grey/dashed lines) or Asbt-deficient (B6.*Slc10a2*^-/-^, solid/colored lines) mice. The 69 reliably detectable species presented here are listed, grouped and color-coded as in Figure S2C. Mean values are from 7 mice per genotype. 1° cBAs, primary conjugated BAs; 1° uBAs, primary unconjugated BAs; 2° cBAs, secondary conjugated BAs; 2° uBAs, secondary unconjugated BAs; 2° metabolites, secondary metabolites; HDPs, hepatic phase 2 detoxification products. **(C)** Similarity matrix of endogenous BA pools in the 4 sites numbered in (A) from wild type or Asbt-deficient mice. Calculated based on levels of the 69 reliably detected BA species described in Figure S2C. Sample IDs list tissue type, genotype and cage number; genotypes and tissues are color-coded at top. **(D)** Heat maps of the relative abundance (Log_2_ fold-change) of individual BA species, determined as above by LC-MS/MS, in the indicated tissues of Asbt-deficient *vs*. wild type mice. Separate heatmaps show BA species found in Figure 1E to be significantly enriched in either wild type siLC (*top*) or feces (*bottom*). \**P*adj < 0.05, Wilcoxon rank-sum test. **(E)** Mean expression (+ SEM; *n* = 7) of BA metabolism-related genes, determined by Nanostring, in whole ileal tissue (*top*) or livers (*bottom*) of wild type (B6) or Asbt-deficient (B6.*Slc10a2*^-/-^) mice. \**P* < 0.05, unpaired two-tailed Student’s *t* test; ns, not significant.

Three responses were observed in Asbt-dependent mice that appeared to reflect coordinated attempts to restore BA homeostasis. First, livers of Asbt-deficient mice increased hepatic BA biosynthesis, presumably to replace 1° cBAs lost to excretion. It is well-established that ileal absorption of 1° cBAs suppresses hepatic BA biosynthesis via two synergistic mechanisms. In one, 1° BAs circulating through ileal enterocytes activate FXR-dependent expression of *Fgf15* (*FGF19* in humans), an enterokine hormone secreted into portal blood that signals through Fgfr4 on hepatocytes to suppress expression of rate-liming enzymes in BA biosynthesis (*i.e*., *Cyp7a1, Cyp8b1*) ^11,17^. In another, 1° BAs returning to the liver via portal blood activate hepatocyte FXR directly, which induces expression of small heterodimer protein (SHP/Nr0b2), and in turn, represses *Cyp7a1*/Cyp*8b1* transcription^15,45,46^. In line with reduced enterohepatic circulation, Asbt-deficient mouse ileal tissue showed lower *Fgf15* expression – despite equivalent *Fxr* expression – while livers of these same mice showed depressed *Shp* and elevated *Cyp7a1*/*Cyp8b1* expression (**Figure 2E**, data not shown).

Second, fecal microbiota from Asbt-deficient mice showed evidence of adapting to increased colonic BA flux by bolstering the relative abundance of organisms carrying BA-metabolizing (BSH, 7α-HSDH) enzymes (**Figure S3A-B**). Enhancement of BA-metabolizing genes within fecal microbial communities of Asbt-deficient mice was explained mostly by a ∼7-fold enrichment of a single unannotated species (CAG-873 sp011959565) carrying both *Bsh* (deconjugation) and 7α-HSDH (dehydroxylation) enzymes; a second *Bsh*- and 7α-HSDH-positive organism, *Muribaculum faecis*, also trended higher in Asbt-deficient communities, although this was not statistically significant (**Figure S3C**). These results were especially notable because wild type and Asbt-deficient mice were reared together in these experiments for months leading up to the day of analysis, to minimize microbial differences and facilitate mixing of enteric microbiomes. Despite this, microbial communities from Asbt-deficient mice were clearly distinct from those in wild type cage-mates, as judged by β-diversity, and showed reduced α-diversity, in line with enrichment of select taxa (**Figure S3D-E**).

Finally, livers of Asbt-deficient mice upregulated BA detoxification pathways, perhaps as a response to increased portal recirculation of microbially-derived 2° uBAs (*e.g*., DCA, LCA). 2° uBAs are more hydrophobic, and thus more cytotoxic at low concentrations, compared with 1° cBAs^4,47,48^. Hepatocytes thus deploy a series of detoxification mechanisms to prevent BA-driven hepatotoxicity, including sulfation and glucuronidation (**Figure S4A**), as well as re-conjugation and basolateral efflux^49,50^. Sulfation is particularly effective at rendering toxic BA species, and other xenobiotics, more hydrophilic and tagging them for urinary or fecal excretion^37,38,51^. DCA and LCA 3-sulfates, but not 24-glucuronides, were increased in Asbt-deficient *vs*. wild type mice (**Figure S4B-C**). BA sulfates were almost entirely restricted to the SI lumen and feces, in line with the concept that BA sulfates are poor substrates for intestinal absorption^52,53^ (**Figure S4C**).

BA sulfation is elicited by binding of 2° BAs to xenobiotic-sensing nuclear receptors (*e.g*., VDR/*Nr1i1*, PXR/*Nr1i2*) in hepatocytes, which in turn promote expression of BA sulfotransferase enzymes encoded by the mouse *Sult2a* gene cluster^48,54,55^ (**Figure S4D**). Once expressed, Sult2a enzymes transfer a sulfonate group to C-3 or C-7 on the BA steroid nucleus from a universal donor, termed phosphoadenosine phosphosulfate (or PAPS), which itself is regenerated by a PAPS-synthase enzyme encoded by the *Papss2* locus (**Figure S4D**) ^51^. Consistent with enhanced hepatic BA sulfation, livers from Asbt-deficient mice displayed a collective increase in *Sult2a* gene expression, but not *Papss2*, compared with wild type controls (**Figure S4E**). Together, these multi-site and multi-omics analyses unmask intricate response networks that follow loss of Asbt-dependent ileal BA reabsorption and underpin BA homeostasis *in vivo*.

### Computational modeling of *in vivo* bile acid metabolism

Endocrine networks that connect the liver, ileum and colonic bacteria, and that coordinately regulate BA homeostasis are difficult and expensive to study through experimentation alone. Thus, we sought to develop mathematical models capable of analyzing BA circulation and metabolism in more integrated and quantitative fashions (**Figure 3A**). A previous model by Sips *et al*. reported that BA transit speed decreases from proximal to distal along the intestinal tract^56-59^, causing BAs to accumulate in the ileum and colon in the absence of intestinal absorption **(Figure S5A)**. However, most BAs are reabsorbed – either actively in the ileum or passively throughout the SI and LI – which drives their eventual reintroduction into the duodenum, together with the pool of newly synthesized BAs from the liver. To monitor and account for each of these features, we incorporated independent rates and locations for active and passive intestinal BA absorption, as well as for hepatic BA synthesis, microbial 2° BA metabolism and BA reintroduction to the duodenum after enterohepatic cycling. Once we established optimized parameters that best fit our experimental data, we analyzed model outputs to determine which aspects of BA circulation and metabolism regulate pool size and complexity.

**Figure 3.**
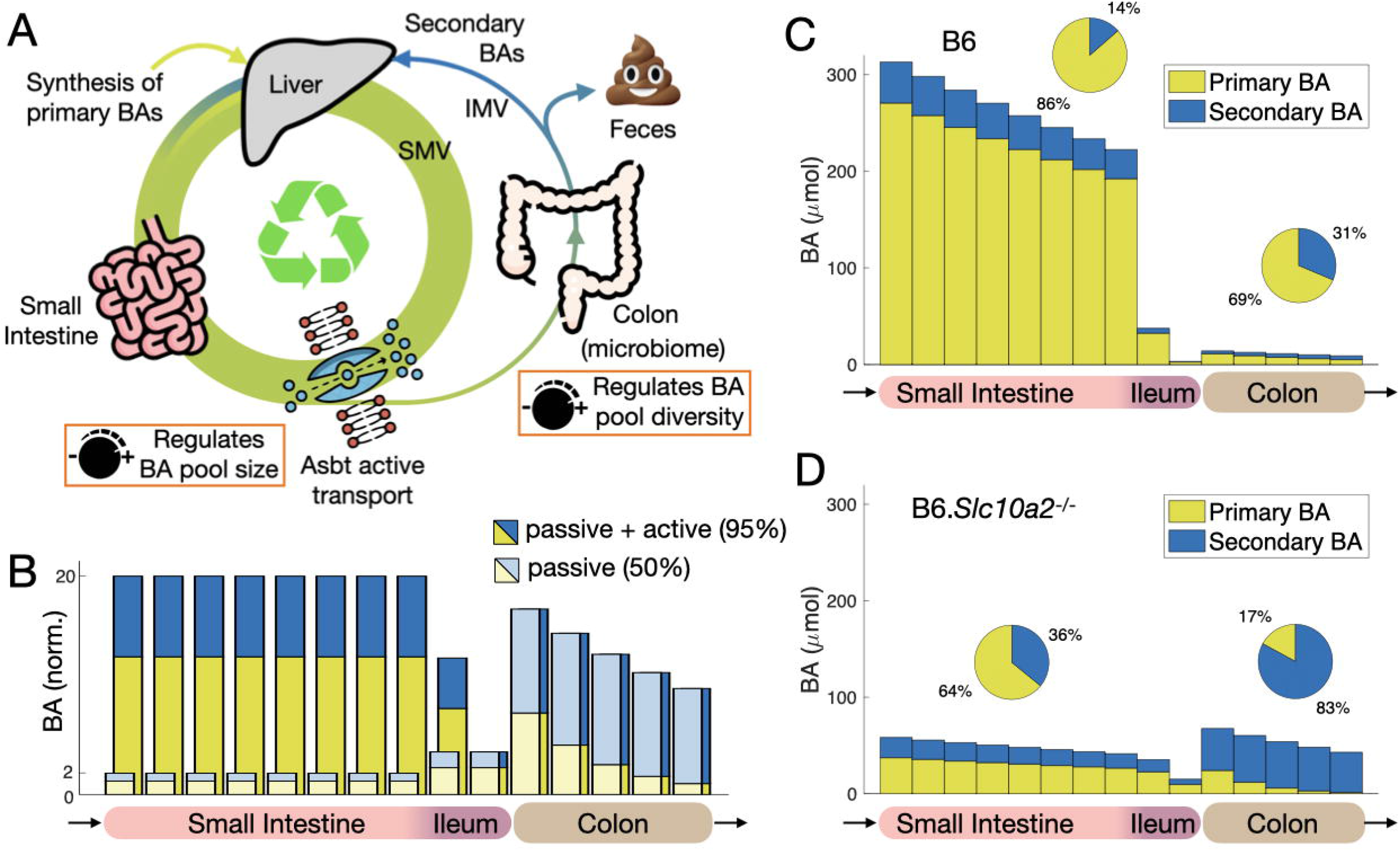
Mathematical modeling of bile acid enterohepatic circulation. **(A)** Schematic diagram of bile acid (BA) enterohepatic circulation. The circulating pool has two inputs: (*i*) 1° BAs synthesized *de novo* by the liver, to replace those not absorbed in the intestines and excrete in feces; and (*ii*) BA that are reintroduced into the small intestine after absorption in the small or large intestines, portal recirculation to the liver, and secretion into bile. This includes 1° BAs that become metabolized into 2° BAs by colonic microbiota and reintroduced by hepatic re-secretion after portal recirculation. IMV, inferior mesenteric vein; SMV, superior mesenteric vein. **(B)** Effect of intestinal absorption of enterohepatic BA pool size and composition. BA levels are scaled by the amount of total BAs in the small intestine (SI) absent intestinal absorption. Combining active absorption in the ileum (90% efficiency) and passive absorption in the colon (50% efficiency) for a total of 95% intestinal absorption increases the small intestinal pool 20-fold relative to no intestinal absorption. Loss of Asbt-mediated active absorption, with only passive absorption remaining, explains the decrease in pool size in B6.*Slc10a2*^-/-^ mice, but not changes in BA pool composition. **(C-D)** Optimization of SI absorption and large intestinal (LI) microbial BA metabolism rates to fit experimental data for C57BL/6J wild type (B6, panel C) and Asbt-deficient (B6.*Slc10a2*^-/-^, panel D) mice. Differences in absorption rates between 1° BA and 2° BA were not considered. Increased conversion of 1° BA into 2° BA by the microbiota in B6.*Slc10a2*^-/-^ mice explains increased BA pool diversity.

At steady state (*i.e*., stable BA circulation), the amount of newly synthesized hepatic BAs introduced into the duodenum matched that which escaped SI absorption and was excreted in feces. When BAs were actively absorbed in the ileum, mimicking wild type mice, this markedly (∼20-fold) increased the size of the BA pool in enterohepatic circulation, and BAs synthesized by the liver to replace those lost to excretion represented only a small fraction (∼5%) of the total pool (**Figure 3B, S5B-C**). As most intestinal BA absorption in our model occurred via active transport in the ileum (**Figure 3C**), only a small fraction of BAs reached the LI, became metabolized to 2° species, and were ultimately reintroduced to the SI pool. This simulation suggested our model was appropriately calibrated to experimental data, and reinforced earlier concepts that Asbt-mediated ileal BA uptake sustains pool size, whereas bacterial BA metabolism in the LI is the major source of pool diversity^60,61^ (**Figure 1C, 2B, 3C**).

Abolishing active BA absorption in our model – to mimic that in Asbt-deficient mice – reproduced the ∼5-fold reduction in SI BA pool size observed in our experiments, but was not sufficient to explain increased fecal BA excretion (**Figure 2B, 3B, S5C**). In this case, the SI BA pool size was reduced markedly, but a far higher fraction of this smaller pool escaped ileal absorption and was excreted. Importantly, increased fecal BA excretion in the absence of active ileal absorption – which are physiologically linked in Asbt-deficient mice (**Figure 2A, 2E**) – only occurred in our model when hepatic synthesis rates were increased (**Figure S5A**). These simulations support and extend our experimental observations in Asbt-deficient mice, where reduced ileal absorption and increased hepatic synthesis may contribute independently to the loss of SI BAs and bloom of fecal BAs, respectively.

Finally, we used our model to interrogate determinants of BA pool complexity. Neither increasing hepatic synthesis nor decreasing active ileal absorption substantially altered representation of bacterially-derived 2° BAs within the SI pool (**Figure 3B, S5A**). In these simulations, increases in hepatic re-secretion of LI-derived 2° BAs were negated by proportionate increases in 1° BA biosynthesis by the liver (**Figure 3B, S5A**). In fact, we tested a range of ileal (and colonic) absorption rates in our simulations, and found no conditions that obviously changed SI BA pool composition (**Figure 3B**). Thus, we considered that the increased proportion of 2° BAs in the SI pool of Asbt-deficient mice might require parallel increases in 2° BA metabolism by LI microbiota (**Figure 3A**). In testing this possibility, we varied LI BA metabolism in our model and searched for rates that best fit our experimental data. These simulations suggested that a 9-fold increase in bacterial BA metabolism rates would be necessary to explain the increased 2° BA representation seen in the SI of Asbt-deficient animals (**Figure 3C-D)**, a number close to the ∼7-fold enrichment of *Bsh*^+^*Bai*^+^ taxa observed experimentally in Asbt-deficient *vs*. wild type fecal microbiomes (**Figure S3B-C**). We attempted to explore the predicted increase in microbial BA metabolism more directly – by assessing metabolism of isotopically labelled (t)CA by cultured fecal microbiota from wild type or Asbt-deficient animals – but these communities, and *bai*^*+*^ taxa in particular, were poorly maintained in our culture conditions (see methods for details). These results illustrate how computational models can enhance experimental studies of *in vivo* BA metabolism, and suggest that hepatic synthesis, ileal absorption, and bacterial metabolism each provide unique inputs to the enterohepatic BA pool.

### Quantifying ileal BA absorption *in vivo*

The estimate that ∼95% of all BAs are absorbed in the ileum stems from classical studies in which small intestines of experimental animals were perfused with individual BAs, or human volunteers were dosed orally, and serum levels were followed over time to estimate absorption^44,62-70^. However, as each BA species possesses unique pharmacokinetic properties, including rates and routes of intestinal absorption^71,72^, this general estimate cannot be uniformly applied to complex BA pools. In addition, neither active nor passive intestinal BA absorption are physiological constants; BA malabsorption is a common manifestation of many intestinal disorders and results from reduced ASBT expression or function, whereas changes in epithelial barrier function – due to genetics, diet or dysbiosis – may affect passive BA absorption^73^. Because intestinal absorption is a major determinant of which luminal BAs ultimately seed mucosal tissues (and at what concentrations), we next asked if our metabolomics dataset (**Figures 1-2**) could be used to infer rates and routes of ileal BA reabsorption *in vivo*.

We first considered that smvB/siLC BA ratios alone might reflect ileal absorption rates. However, these were similar between wild type and Asbt-deficient mice, both for individual BA species [*e.g*., (t)βMCA] and for the pool as a whole (**Figure S6A**). In considering this result further, we realized that such a ratio assumes siLC and smvB BA levels are maintained in a constant state of disequilibrium in the absence of Asbt (*i.e*., that only smvB, but not siLC, BAs decrease in Asbt-deficient mice; **Figure S6B, *left***). To the contrary, our experimental data and computational modeling better supported a model in which both siLC and smvB BAs are depleted in Asbt-deficient mice, and where the BAs remaining in enterohepatic tissues circulate at a new equilibrium limited by passive intestinal absorption rates. (**Figure 2A, 4A, S6B, *right***). In exploring relationships between siLC and smvB BA levels further, we found that the natural variation in siLC and smvB BA levels across cohorts of wild type or Asbt-deficient mice generated line slopes (*m*), which were generally lower in Asbt-deficient *vs*. wild type animals, and which better intimated absorption rates. By defining these slopes (absorption rates) for each BA species, and in both wild type and Asbt-deficient animals, we were able to use standard line equations (y = *m*x + b) to estimate amounts of active and passive absorption, both for individual BA species and the total pool (**Figure 4B, 4C, *top***).

**Figure 4.**
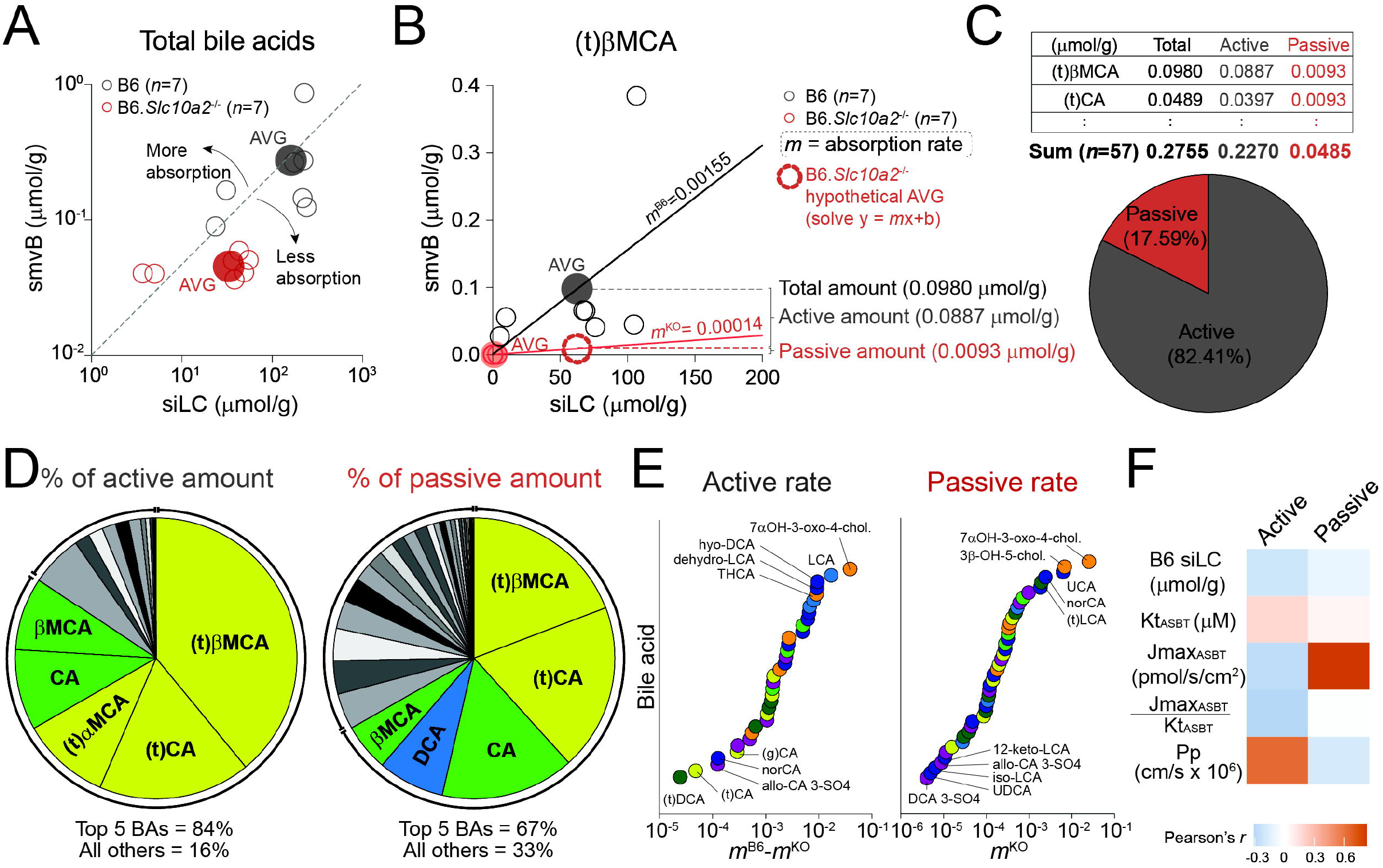
Estimating active and passive ileal bile acid reabsorption from *ex vivo* metabolomics data. **(A)** Relationship between total bile acid (BA) concentrations in small intestine luminal content (siLC, x axis) and superior mesenteric vein blood (smvB, y axis) of the same C57BL/6J wild type (B6, grey) or Asbt-deficient (B6.*Slc10a2*^-/-^, red) mice (*n* = 7). Sum totals of 69 reliably detectable BA species (described in Figure S2B-C) for individual mice are shown in small, clear circles; averages values per genotype are indicated by large, filled circles. Dashed diagonal line shows the y=x axis. **(B)** Example of how active and passive ileal reabsorption is calculated for individual BA species. Individual (small, clear circles) and average (large, filled circles) siLC and smvB concentrations of (t)βMCA in wild type (grey) or Asbt-deficient (red) mice are shown (*n* = 7). Linear regression analyses, per genotype, produced line slopes (*m*) indicative of ileal absorption rates. Line equations (y = mx+b) were used to determine the amount of each BA – if present in Asbt-deficient mouse siLC at the same concentration as in wild type mouse siLC (x) – would be absorbed and enter smvB (y). Since the amount in wild type smvB (large, filled grey circle) is known, and represents total ileal absorption when both active and passive ileal absorption are intact, and we solved for amount of passive BA absorption in Asbt-deficient smvB (large, clear red circle), where only passive BA absorption occurs, the difference between wild type and Asb-deficient y-axis points equals the amount of active (Asbt-mediated) ileal absorption. **(C)** *Top*, amount of total, active and passive ileal absorption (in μmol/g) inferred for 57 individual BAs, as in (B). Amounts for (t)βMCA and (t)CA are shown as examples. Absorption amounts for all 57 BA species are shown in Figure S6C. *Bottom*, pie chart showing the percentage of total ileal BA absorption, calculated as in (B) and Figure S6C, attributed to active (Asbt-mediated, grey) or passive (red) absorption. **(D)** Pie charts showing percentages of total active (*left*) or passive (*right*) ileal BA absorption, calculated as in (B-C), for each BA species. Total percentages of the top 5 BAs *vs*. all others are listed at bottom. **(E)** Rank-orders of all 57 BAs from lowest to highest active (*left*) or passive (*right*) ileal absorption rates (*m*), calculated as in (B) and color-coded based on putative biosynthetic origin as in Figure S2C. **(F)** Heatmap of Pearson correlation (*r*) values between rates (*m*) of active or passive ileal absorption rates, inferred here [as in (B, E)], with previously-measured physicochemical properties for 10 major primary and secondary BAs^71^. Kt, Asbt transport constant; Jmax, maximal amount of Asbt-dependent transport (both theoretically related to active, Asbt-mediated BA transport); Pp, passive permeability (theoretically related to passive BA absorption).

After removing 12 additional BAs whose median concentration in wild type siLC was 0, we calculated the size of the ileal BA pool in fasted wild type mice to be 0.2755 μmol/g (**Figure 4C, S6C**). More than 80% of this pool could be attributed to Asbt-dependent transport, and ∼85% of actively-transported BAs was comprised of only five abundant 1° BA species, including conjugated and unconjugated CA, as well as the rodent-biased muricholates [*e.g*., (t)αMCA, (t)βMCA, βMCA; **Figure 4D**]. 1° cBAs, which are thought to strictly require active absorption for their enterohepatic circulation^71^, also displayed surprisingly high levels of passive absorption – presumably due to sheer abundance – whereas passive absorption also facilitated ileal entry of many less abundant, hydrophobic secondary BAs (*e.g*., DCA), thus increasing diversity within the absorbed pool (**Figure 4D**). Intriguingly, our inferred rates of *in vivo* active ileal absorption (*m*^B6^ - *m*^KO^) did not correlate with previously determined Asbt transport kinetic parameters (*J*_max_, *K*_T_) for 10 individual BAs using standard *in vitro* MDCK cell-based transport assays^71^ (**Figure 4E-F**). Similarly, *in vivo* passive absorption rates (*m*^KO^) failed to correspond to *in vitro*-defined passive permeability (*P*_P_) estimates from the same *in vitro* study^71^ (**Figure 4E-F**). Whether these discrepancies speak more to competitive dynamics between individual BAs within complex pools, or to other *in vivo* variables (*e.g*., BA retention within mucosal tissue, etc.), remains unclear. Regardless, these results provide a framework for understanding ileal BA pool dynamics in mice and represent an important next step in relating putative BA functions to *in situ* concentrations.

### Functional consequences of aberrant BA metabolism in Asbt-deficient mice

Much of contemporary BA research (*i.e*., since the discovery of FXR as an endogenous BA receptor 25 years ago^12-14,74^) has centered on defining interactions with host receptors, and elucidating how such interactions impact GI physiology. Yet with few exceptions, BA functional studies have focused on individual species – not complex pools – leaving considerable gaps in knowledge of how endogenous BAs influence host intestinal function *in vivo*. We sought to leverage our newly acquired insights into the size and complexity of mouse ileal BA pools (**Figure 4**) to explore their cellular and molecular functions.

After extracting bulk metabolites from siLC of fasted and co-housed wild type or Asbt-deficient mice, which included but were not limited to BAs, contents from each animal were divided, and either left intact or depleted of BAs using a BA-binding resin, cholestyramine (CME), (**Figure 5A**). Total BA concentrations were then normalized between the two intact (wild type, Asbt-deficient) extracts – to focus downstream analyses on compositional differences – and individual species were quantified by LC-MS/MS. After accounting for dilution factors used in functional studies (below), which were designed intentionally to approximate the size of the ileal BA pool in fasted wild type mice (0.2755 μmol/g, ∼ 291 μM; **Figure 4C, S6C**), we confirmed that the mean total BA concentration in intact wild type siLC extracts was ∼286 μM; CME treatment efficiently removed ∼97% of these BAs (**Figure 5A**). Intact extracts from Asbt-deficient mice contained ∼2-fold fewer total BAs (mean total concentration ∼112 μM), and CME again depleted ∼93% of these (**Figure 5A**). At the species level, CME treatment depleted 55 of 69 reliably detected BAs by at least 95% in both wild type and Asbt-deficient siLC extracts; only 2 minor BA metabolites (7-keto-DCA, UCA) were depleted with <75% efficiency (**Figure 5B**). Most importantly, intact siLC extracts produced here displayed the same Asbt-dependent hallmarks as before (**Figure 2**) – reductions in 1° cBAs, and increases in 1° uBAs, 2° uBAs and BA 3-sulfates, in Asbt-deficient *vs*. wild type extracts (**Figure 5A**).

**Figure 5.**
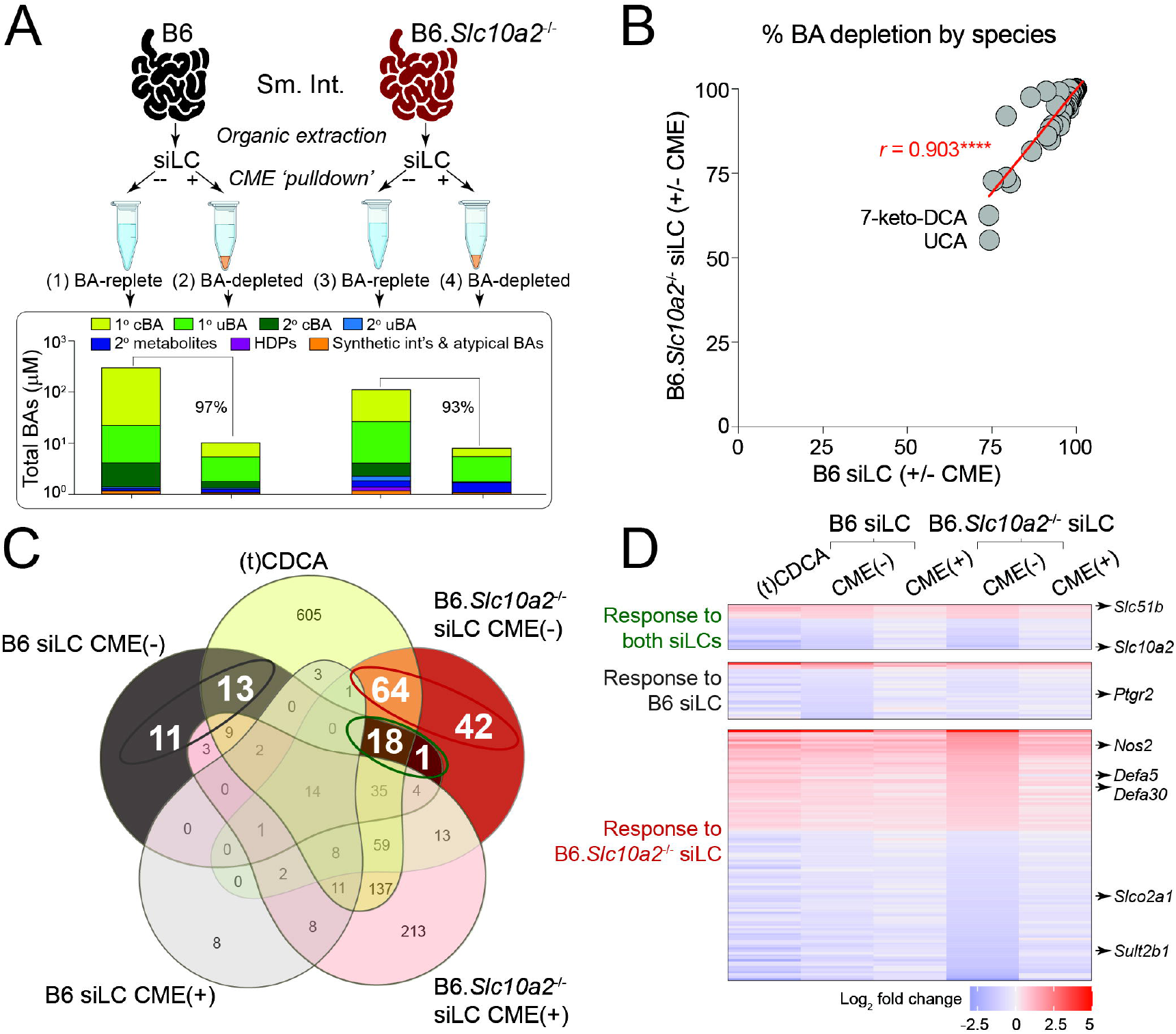
Compositionally distinct small intestinal bile acid pools induce divergent transcriptional responses in cultured ileal explants. **(A)** *Top*, schematic diagram of the experimental workflow for interrogating direct, *ex vivo* functions of endogenous, enterohepatic bile acid (BA) pools in mice. Organic extracts from small intestine luminal content (siLC) of C57BL/6J wild type (B6, black) or Asbt-deficient (B6.*Slc10a2*^-/-^, red) mice were divided, and either treated with cholestyramine (CME) to deplete BAs (BA-depleted), or left intact (BA-replete). *Bottom*, mean total BA concentrations (*n* = 3), determined by LC-MS/MS, in siLC extracts indicated above (BA-replete or BA-depleted from wild type or Asbt-deficient mice). Proportions of discrete BAs in defined categories are color-coded as in Figure 1C, 2B, S2C. **(B)** Efficiency of CME-mediated BA-depletion in siLC extracts from C57BL/6J wild type (B6) or Asbt-deficient (B6.*Slc10a2*^-/-^) mice. Each symbol represents an individual BA species; values were calculated based on mean BA concentrations (μmol/g) in CME-treated *vs*. untreated siLC extracts [100 – (+CME concentration/-CME concentration)] from 3 mice per genotype. Pearson correlation (*r*) value, in red, demonstrates nearly identical BA depletion efficiencies by CME pre-treatment in both wild type and Asbt-deficient siLC extracts. \*\*\*\**P* < 0.0001, Pearson correlation test. **(C-D)** Venn diagrams of gene numbers (C) and heatmap of gene expression (D) for genes significantly altered (*P*adj < 0.05, DESeq2; as in Figure S7E, *n* = 3-4) in cultured ileal explants 8 hr after treatment with 200 μM (t)CDCA, or equal dilutions of intact (CME-) or BA-depleted (CME+) siLC extracts from wild type (B6) or Asbt-deficient (B6.*Slc10a2*^-/-^) mice. Intact extracts were normalized to produce 300 μM total BAs in each explant cultured, based on colorimetric kit analysis as in Figure S7D; the same dilutions of matched BA-depleted extracts were used. Data were pooled from 2-independent experiments (1 or 3 biological replicates in each), using siLC extracts from separate wild type or Asbt-deficient mice.

To assess functions of these distinctive BA pools, we employed intestinal explant cultures^75^. In optimizing culture conditions, we found that wild type mouse ileal, but not colonic, explants responded rapidly (within 4-8 hr) to low micromolar concentrations of (t)CDCA by upregulating the canonical FXR target gene, *Fgf15*; similar results were seen with the synthetic FXR agonist, GW4064^76^ (**Figure S7A-B**). Blocking Asbt transport function with the FDA-approved small molecule, odevixibat^77^, blunted (t)CDCA-induced *Fgf15* expression in ileal explants (**Figure S7C**), confirming that FXR activation by conjugated BAs in ileal explants – as *in vivo* – requires Asbt-mediated uptake by enterocytes. Intact siLC extracts from wild type mice also evoked *Fgf15* upregulation, akin to (t)CDCA alone, whereas this response was muted in the presence of parallel BA-depleted extracts (**Figure S7D**).

Having established the system, we went on to use RNA-seq to compare genome-wide transcriptional responses of wild type ileal explants to (t)CDCA alone, or to the four SI extracts generated above (*i.e*., from wild type or Asbt-deficient mice with or without BA depletion). To quantify similarities and differences between ileal explant responses to wild type or Asbt-deficient BA extracts, we enumerated genes whose expression was significantly affected (*i.e*., induced, repressed) by each BA extract *vs*. vehicle (DMSO) alone (**Figure S7E**). Importantly, fewer than 150 genes displayed differential expression in the presence of either or both BA-replete, but not BA-depleted, siLC extracts, in contrast to more than 900 genes that were influenced by a typical (supraphysiologic) concentration of (t)CDCA (200 μM; **Figure 5C, S7E**). On its own, this result reveals unique layers of information that can be extracted upon examining host tissue responses to physiologically-defined BA pools.

Of the nearly 150 genes regulated by either or both BA-replete, but neither BA-depleted, siLC extracts, a large majority (*n*=130; 87%) reflected unique responses to wild type or Asbt-deficient BA pools (**Figure 5C-D**). Asbt-deficient extracts elicited more unique transcriptional responses in ileal explants (*n*=106) than those evoked by wild type extracts (*n*=24) (**Figure 5C, S7E**). Unique responses of ileal explants to the two SI BA pools spanned multiple key areas of intestinal physiology, from prostaglandin (*Ptgr2*) and nitric oxide (*Nos2*) metabolism, to antimicrobial activity (*Defa5, Defa30*) and xenobiotic metabolism (*Slco2a1, Sult2b1*) (**Figure 5D**). Just over half (∼52%) of each uniquely responsive gene set (*i.e*., to wild type or Asbt-deficient BA-replete extracts) were predicted by the explant response to (t)CDCA alone, whereas nearly all (18/19, 95%) of the genes commonly influenced by both wild type and Asbt-deficient BA pools – including FXR gene targets *Fgf15, Slc51b* (OSTβ) and *Slc10a2* (Asbt) – were similarly affected by (t)CDCA (**Figure 5C-D, S7F**). These results establish new paradigms in the functions of endogenous BA pools and illustrate how aberrant BA metabolism can directly impinge on host intestinal physiology.

## Discussion

Appreciation continues to grow for the complex endocrine connections between the liver, intestines and gut microbial communities that underpin most aspects of GI physiology, including the circulation and metabolism of BAs. Nevertheless, most contemporary human or mouse BA studies continue to use single-site measurements, most commonly in plasma (to catalog BAs as biomarkers), liver/gallbladder (to explore BA biosynthetic mechanisms), ileum (to decipher BA absorptive and signaling mechanisms) or stool (to interrogate microbial metabolic activities). The results presented here suggest that BAs, gut microbial communities and host enterohepatic tissues should be viewed as integrated functional units; perturbation of any one node necessarily affects the others, and confounds experimental readouts if all nodes are not monitored and controlled for in parallel. The approaches, analyses and models presented here were designed to serve as an integrated methods set, such that microbiome scientists can monitor effects of dysbiosis on hepatic metabolism or intestinal absorption, and vice versa.

Our multi-site BA analysis quantifies the striking compartmentalization of SI and LI BA pools. Although this result was largely anticipated, based on historical knowledge and more recent large-scale metabolomics studies^78^, strict quantitative comparisons described here have important implications for multiple areas of biology. Compartmentalization of SI and LI BA pools stands to influence everything from the behavior of local immune and epithelial subsets (discussed below), to that of small *vs*. large bowel microbes and cancers. Compartmentalization also suggests that analysis of fecal BAs alone – an increasingly common measurement in mice and humans – has limited relevance to features in enterohepatic circulation. This is further complicated by the result that BAs in peripheral circulation are even more diverse and variable than those in feces, displaying mixed features of both SI and LI BA pools. Thus, great care must be taken when interpreting fecal or serum BA data alone in patients and controls.

Our results offer working answers to long-standing questions in BA research. For example, by parallel sampling of BA pools in siLC and smv of individual mice, we were able to estimate the size, complexity and routes of absorption of the ileal BA pool in mice. Our results suggest that, of the ∼95% of BAs absorbed in the wild type mouse ileum at steady-state, more than 80% of this pool can be attributed directly to Asbt-dependent transport, whereas less than 20% is accounted for by passive (Asbt-independent) diffusion. As expected, the vast majority (∼70%) of ileal BAs in fasted wild type mice were comprised of a select set of the most abundant primary conjugated and unconjugated BAs in mice, namely CA and α/βMCA. At the same time, we observed surprisingly high passive absorption of taurine-conjugated primary BAs *in vivo*, which are generally viewed as requiring Asbt-mediated transport for ileal entry, given relatively low passive permeability^71^. Intriguingly, neither active nor passive *in vivo* ileal absorption rates inferred here were predicted by prior *in vitro* studies in which individual primary or secondary BAs were individually added to MDCK cells with or without Asbt expression^40^. One potential explanation for this discrepancy is *in vivo* competition between myriad unique BA species within intestinal pools for finite numbers of (Asbt) transport proteins. If true, *in vitro* Asbt transport kinetics of individual BAs, such as (t)CA, should be influenced by the presence of other species. It also seems plausible that more complex *in vivo* variables, for example retention of BAs in intestinal tissues and/or egress from tissues into portal blood, inherently limit the resolution with which one can interrogate Asbt transport kinetics *in vivo*. Finally, it is worth noting that we used linear regression analyses to estimate ileal absorption rates *in vivo* (**Figure 4A-B**), whereas Asbt transport kinetics may be better approached using semi-logarithmic curve-fitting to follow Michaelis–Menten enzyme principles. Whatever the underlying dynamics, the properties of the mouse ileal BA pool proposed here can refined in future studies, but also expanded to understand whether or how ileal BA absorption changes throughout the post-prandial period, in response to different diets or disease models, or across discrete inbred mouse strains (*e.g*., C57BL/6, BALB/C, FVB/N).

Through combined *in vivo* experimentation and mathematical modeling, we show that Asbt-mediated ileal BA absorption not only maintains the size of the SI BA pool; it also safeguards SI pool composition, particularly the enrichment of 1° cBAs. At steady-state, the SI BA pool is enriched in 1° cBAs and specialized for lipid and fat-soluble vitamin absorption. Conversely, loss of Asbt-mediated absorption not only reduced the size of the enterohepatic BA pool ∼5-fold; it also became enriched in hydrophobic 2° BAs generated by bacterial metabolism in the LI. Mechanistically, our results suggest that infiltration of 2° BAs into the SI pool likely involves enhanced 2° BA metabolism by colonic bacteria. However, we cannot exclude the possibility that enrichment of 2° BAs within the SI pool of Asbt-deficient mice also entails preferential active transport of 1° BAs and/or biased hepatic synthesis of CA versus MCA in Asbt-deficient settings, which were not accounted for in our model. Regardless, marked compositional shifts in the SI BA pool of Asbt-deficient mice raise intriguing clinical questions that remain unresolved. For example, our results suggest that the composition of enterohepatic BA pools may be overtly changed in individuals with intestinal diseases, surgical procedures, or pharmacological interventions that decrease ASBT expression or function. If so, do these pools display disproportionately high levels of 2° BAs? And does this enrichment of 2° BAs affect metabolic or nutritional properties of the enterohepatic pool, such as facilitation of lipid and fat-soluble vitamin absorption? These and other questions will be critical to address to ultimately achieve broader perspectives of how BA metabolism impacts human health and disease.

Our modeling results also predict that ileal BA absorption and hepatic BA biosynthesis act independently to determine the size of the SI BA pool and amount of fecal BA excretion, respectively. Although seemingly incremental, this under-appreciated principle provides a mechanistic explanation for the reported uncoupling of ileal BA absorption and fecal BA excretion in mice or humans lacking OSTα/β-mediated basolateral BA efflux in ileal enterocytes. Despite a block in ileal BA absorption, fecal BA excretion is not elevated in OSTα/β-deficient pre-clinical or clinical settings because of increased gut-liver signaling via Fgf-15 (in *Slc51a*^-/-^ mice) or FGF-19 (in children born with homozygous loss of *SLC51A* function), which act in a dominant fashion to prevent increases in hepatic BA biosynthesis despite reduced enterohepatic BA circulation^79,80^.

Finally, our study establishes new paradigms regarding functions of endogenous BA pools. Many studies to-date have explored molecular functions of individual BAs, either *in vitro* or *in vivo*. Yet *in vivo* studies – for example where mice are fed BA-supplemented diets^19,28^ – are confounded by effects on host (*e.g*., hepatic) and gut microbial metabolism (**Figure 2-3, S3-4**), whereas *in vitro* studies are limited by routine use of supraphysiologic BA concentrations and do not account for *in vivo* competitive dynamics. The concept of BA competition becomes especially important when one considers that discrete BA species have been shown to bind competitively to the same NR, but elicit opposite functions – CDCA activates FXR, whereas α/βMCA suppress it^74,81^. Still, the extent to which these or other competitive interactions shape host intestinal responses to BAs have yet to be rigorously examined.

Using cultured ileal explants, we show that direct *ex vivo* functions of two SI BA pools with markedly different compositions – in this case from wild type or Asbt-deficient mice – are more different than they are similar. Despite possessing ∼2-fold fewer total BAs after normalization attempts, SI BA extracts from Asbt-deficient mice nonetheless elicited more differential gene expression that those evoked by wild type BA pools. Although further studies are needed to fully elucidate the molecular bases for these differential signaling activities, our results are consistent with the possibility that different ratios of FXR-activating (*e.g*., CDCAs) *vs*. - inhibitory (*e.g*., α/βMCAs) BAs exist within the pools. Both CDCAs and α/βMCAs were substantially less abundant in Asbt-deficient *vs*. wild type SI extracts (**Figure 2D**), but loss of Asbt disproportionately depleted muricholate species, resulting in a ∼20-fold relative increase in the CDCA:MCA ratio. On the other hand, several known FXR target genes – *Fgf15, Slc51b* (OSTβ) and *Slc10a2* (Asbt) – were similarly influenced by both wild type and Asbt-deficient SI BA pools, suggesting that differential FXR activation is not the only source that underlies unique transcriptional responses of ileal explants to the two BA pools. Another, likely additive, signaling activity enriched within the Asbt-deficient SI BA pool is the increases in multiple 2° BA species and metabolites – including DCA, LCA and their corresponding metabolites – known for activating orthogonal NR pathways, such as VDR and PXR^48,82^.

Ultimately, substantial work remains to fully appreciate how BA dysmetabolism impacts host physiology, and in turn, to leverage these insights for therapeutic benefit in patients. However, this study takes key next steps by empirically defining the form and functions of ileal BA pools in mice. These approaches, when combined, will be broadly useful for interrogating functions of BA pools related to age, sex, disease, or treatment differences. In addition, the paradigms established here are forward-looking and should inform the metabolism, intestinal absorption and functions of recently, or still yet-to-be identified. BAs.

## Acknowledgements

The authors thank Core Facility staff at UF Scripps, the Genomics and Molecular Biology Shared Resources at Dartmouth’s Norris Cotton Cancer Center with NCI Cancer Center Support Grant 5P30 CA023108-37, and Creative Proteomics for technical support. We further acknowledge Drs. Casey Weaver (University of Alabama-Birmingham) and Maria Abreu (University of Miami) for critical discussions. Financial support was from National Institutes of Health (NIH) grants ES 033988-01A1 (to G.A.O.), T32HL134598 (to K.E.B.), DK047987 (to P.A.D.), and R01AI143821, R01AI164772 and U01AI163063 (to M.S.S.), as well as a grant from the Canadian Institutes of Health Research (PJT-186117; to H.M.K.).

## Author contributions

Conceptualization: P.A.D., D.S., M.S.S. Software: K.S., A.D.E., S.S., J.L., A.B. H.D.D. Validation & Investigation: K.S., A.D.E., S.S., K.E.B, A.B., C.J.S., M.J.J., C.L.H., D.S. Formal analysis: K.S., A.D.E., S.S., K.E.B., A.B., H.D.D., D.S., M.S.S. Writing - Original Draft: K.S., D.S., M.S.S. Writing - Review & Editing: K.S., A.D.E., J.L., A.R.P., G.A.O, S.J.K., P.A.D., D.S., M.S.S.

## Supplemental information

Supplemental information for this manuscript includes supplemental figures (Figure S1-S9), one supplemental table (Table S1) and supplemental methods.

### Figure legends

**Figure S1. Representative LC-MS/MS chromatogram of bile acid analysis (related to Figure 1-2, 4-5).** Mass/charge (m/z) peaks for exemplar bile acid (BA) species in a representative C57BL/6J wild type mouse small intestine luminal content (siLC) sample are annotated.

**Figure S2. Pre-processing, filtering and clustering of mouse *ex vivo* bile acid metabolomics data (related to Figure 1-2, 4-5). (A)** Concentrations (μmol/g) of individual bile acid (BA) species, determined by LC-MS/MS as in Figure S1, in feces or colon luminal content (cLC) of separate 30 wk-old wild type C57BL/6J (B6) mice (*n* = 1). Pearson correlation shows significant compositional similarity between the BA pools, despite them being from independent animals.

**(B)** Median concentrations (μmol/g; *n* = 7) of individual BA species in the indicated sites of 30 wk-old wild type B6 mice. siLC, small intestine luminal content; smvB, superior mesenteric vein blood; pB, peripheral blood. 88 total BA species were analyzed in each site. Numbers of BA species with median values of >0 in each site are indicated on x-axis; white boxes (*i, ii, iii*) highlight BA species whose median values were 0 in all three gastrointestinal (GI) sites and were removed from downstream analyses. None of the BA species with median values of 0 in pB were excluded, as each of these were consistently detected in at least one of other 3 sites (siLC, smvB, feces).

**(C)** Grouping of the 88 individual BA species analyzed based on: (1) reliability of detection by LC-MS/MS [all except those where median concentrations *i + ii + iii* = 0 in (B)]; and (2) likely biosynthetic origin. 1° cBAs, primary conjugated BAs; 1° uBAs, primary unconjugated BAs; 2° cBAs, secondary conjugated BAs; 2° uBAs, secondary unconjugated BAs; 2° metabolites, secondary metabolites; HDPs, hepatic phase 2 detoxification products. Poorly detected species are not strictly grouped by presumed biosynthetic origin.

**(D)** Violin plots of α-diversity (Shannon index; *n* = 7) within indicated BA pools of wild type B6 mice analyzed by LC-MS/MS as above. \*\**P* < 0.01, Friedman test with Dunn correction for multiple comparisons; ns, not significant.

**(E)** Circos plot of correlations among the 69 reliably detected BA species in all four sites of wild type B6 mice. Line colors represent positive (red) and negative (blue) Pearson correlation values; only lines with < -0.99 or > 0.99 correlation values are shown.

**Figure S3. Enrichment of bile acid-metabolizing taxa in Asbt-deficient mouse gut microbiota (related to Figure 2). (A)** Schematic diagram of genes and chemical reactions involved in bacterial bile acid (BA) metabolism, adapted from^83^. Bai operon, bile acid-induced operon; BSH, bile salt hydrolase; (t)CA, tauro-cholic acid; DCA, deoxycholic acid.

**(B)** Relative abundance of 34 metagenome-assembled genomes (MAGs) observed in feces from 7 pairs of C57BL/6J wild type (B6) or Asbt-deficient (B6.*Slc10a2*^-/-^) mice. Mice were co-housed at weaning (3-to 4-wk of age) and reared together until 30 wk of age. Fresh fecal pellets were collected from individual mice after separation and fasting for 4 hr. Individual taxa are colored and annotated based on presence of *Bsh* and/or *7α-hydroxysteroid dehydrogenase* (*7α-hsdh*) genes; *Bsh* alone (purple), *Bsh* and *7α-hsdh* (yellow), or neither (teal).

**(B)** Relative abundance of *Bsh*- and *7α-hsdh*-positive taxa (CAG-873 sp011959565, CAG-873, *left*; *Muribaculum faecis*, M. faecis, *right*), as determined in (B), within fecal microbiota of wild type or Asbt-deficient mice (*n* = 7). \**P* < 0.05, Wilcoxon signed-rank test.

**Figure S4. Increased hepatic bile acid sulfation in Asbt-deficient mice (related to Figure 2). (A)** Schematic diagram of enterohepatic bile acid (BA) circulation adapted from^59^. BAs recirculating from the small or large intestines can be modified in hepatocytes via sulfation or glucuronidation. DCA, deoxycholic acid; LCA, lithocholic acid; GB, gallbladder; HPV, hepatic portal vein; imv, inferior mesenteric vein; IVC, inferior vena cava; pB, peripheral blood; siLC, small intestine luminal contents; smvB, superior mesenteric vein blood.

**(B)** Concentrations (μmol/g; *n* = 7) of secondary BAs – DCA (*top*) or LCA (*bottom*) – in smvB of C57BL/6J wild type (B6) or Asbt-deficient (B6.*Slc10a2*^-/-^) mice. Horizontal lines indicate means; \**P* < 0.05, paired two-tailed student’s *t* test.

**(C)** Concentrations (μmol/g; *n* = 7) of secondary BA phase 2 detoxification products – DCA (*top*) or LCA (*bottom*) 3-sulfates (3-SO4) or 24-glucuronides (24-gluc) – in the indicated sites of wild type or Asbt-deficient mice. Horizontal lines indicate means; \**P* < 0.05, paired two-tailed student’s *t* test (between the same sites of wild type or Asbt-deficient mice); ns, not significant.

**(D)** Schematic diagram of the BA 3-sulfation pathway. BA sulfotransferase enzymes, encoded by the *Sult2a* gene cluster in mice, transfer a sulfonate (*a.k.a*., sulfate) group (^-^O_3_SO, highlighted red) from the universal sulfonate donor, phosphoadenosine phosphosulfate (PAPS) to C-7 of the BA steroid nucleus. This produces phosphoadenosine phosphate (PAP) as a byproduct, which is reconverted to PAPS by the PAPS-synthase enzyme, *Papss2*.

**(E)** Mean expression (+ SEM; *n* = 7) of *Sult2a* and *Papss2* mRNA, determined by Nanostring, in livers of the same co-housed wild type (B6) or Asbt-deficient (B6.*Slc10a2*^-/-^) mice analyzed above. \**P* < 0.05, unpaired two-tailed Student’s *t* test; ns, not significant.

**Figure S5. Effects of intestinal reabsorption on bile acid pool size and composition (related to Figure 3). (A)** Changes of bile acid (BA) synthetic rates in the liver were considered in the model. Total amount of BAs along the intestinal tract in the absence of reabsorption was shown. Two different rates of hepatic BA synthesis (synthesis=1, lower; synthesis=2, higher) were used. BAs accumulate in the ileum and colon because of slower transit. The higher BA synthetic rate scales up BA amounts along the intestinal tract.

**(B)** Passive BA reabsorption in the colon was considered. In this simulation, a fraction of the BA was recycled, increasing the pool size, and introducing 2° BA into the circulating pool in the SI. A 50% reabsorption efficiency doubles the size of the BA pool in the SI.

**(C)** Active BA reabsorption in the ileum was considered. This model (assuming 90% ileal absorption efficiency) increased the size of BA pool 20-fold, compared with the absence of ileal absorption; note the size of the colonic BA pool does not change. Without passive absorption in the colon, there are no secondary BA introduced in the circulating pool.

**Figure S6. Inferring ileal bile acid absorption from bile acid levels in the small intestinal lumen and superior mesenteric vein (related to Figure 4). (A)** Ratios (+ SEM; *n* = 7) of superior mesenteric vein blood (smvB)/small intestine luminal content (siLC) bile acid (BA) concentrations, determined by LC-MS/MS, in C57BL/6J wild type (B6) or Asbt-deficient (B6.*Slc10a2*^-/-^) mice. Ratios are shown for both the primary conjugated mouse BA, tauro-beta-muricholic acid, (t)βMCA; *left*) and for the sum of all BAs (total BAs, *right*); ns, not significant, Wilcoxon signed-rank test.

**(B)** Schematic diagrams representing two distinct models for how loss of Asbt-mediated ileal BA transport affects enterohepatic BA circulation. Option 1 (at left) assumes disequilibrium between siLC and smvB BA levels in the absence of Asbt. Here, 95% of all BAs in siLC of wild type mice (in B6 mice, *top*) are reabsorbed in the ileum and return to the liver via smvB. Loss of Asbt (in B6.*Slc10a2*^-/-^ mice, *bottom*) reduces the amount of smvB BAs (in this example, from 95% of siLC BAs in wild type mice to 20% of siLC BAs in Asbt-deficient mice), but siLC BA levels remain unchanged. Option 2 (at right) assumes that both siLC and smvB BAs are depleted in the absence of Asbt.

**(C)** Amount of active, passive and total ileal absorptions for each BA species, calculated as in Figure 4B-C.

**Figure S7. Transcriptional responses of cultured ileal explants to synthetic and endogenous bile acids (related to Figure 5). (A)** *Left*, kinetics of *Fgf15* mRNA upregulation in cultured ileal explants after treatment with 200 μM tauro-chenodeoxycholic acid [(t)CDCA] or vehicle (DMSO) alone. *Right*, dose response of (t)CDCA on *Fgf15* mRNA expression in cultured ileal explants after 8 hr. Data are presented as mean RQ values (+ SEM), determined by qPCR, in 3-independent experiments using ileal explants from separate C57BL/6J wild type (B6) mice; average values from 3 technical replicates (*i.e*., 3 pieces of ileal tissue) were calculated in each independent experiment. \*\**P* < 0.01, two-way ANOVA with Sidak correction for multiple comparisons.

**(B)** Mean *Fgf15* mRNA expression (+ SEM; *n* = 9) in cultured ileal (*left*) or proximal colon (prox. colon, *right*) explants, determined by qPCR as in (A), after treatment with or without 200 μM (t)CDCA or 200 μM GW4064 (FXR agonist) for 8 hr. \**P* < 0.05, \*\*\*\**P* < 0.0001, one-way ANOVA with Dunnett correction for multiple comparisons; ns, not significant.

**(C)** Mean *Fgf15* mRNA expression (+ SEM; *n* = 8-9) in cultured ileal explants, determined by qPCR as in (A-B), after treatment with 200 μM (t)CDCA or 200 μM GW4064 (FXR agonist) in the presence or absence of the Asbt inhibitor, odevixibat (10 μM) for 8 hr. \**P* < 0.05, unpaired two-tailed Student’s *t* test.

**(D)** *Left*, total bile acid (BA) concentrations (+ SEM; *n* = 7), presented as μmol/g and determined by colorimetric kit, in C57BL/6J wild type (B6) small intestine luminal content (siLC) extracts with or without cholestyramine (CME)-mediated BA depletion. *Right*, Mean *Fgf15* mRNA expression (+ SEM; *n* = 7-9) in cultured ileal explants, determined by qPCR as in (A-C), after 8 hr treatment with or without 200 μM (t)CDCA, or equivalent dilutions of intact or BA-depleted siLC extracts normalized to add 200 μM total BAs in intact extracts. \*\**P* < 0.01, \*\*\**P* < 0.001, Kruskal-Wallis test with Dunn correction for multiple comparisons or Wilcoxon signed-rank test.

**(E)** Differential gene expression (*P*adj < 0.05), determined by DESeq2 analysis of RNA-seq data (*n* = 3-4), in cultured ileal explants treated for 8 hr with 200 μM (t)CDCA, or equal dilutions of intact (CME-) or BA-depleted (CME+) siLC extracts from wild type (WT; *i.e*., B6) or Asbt-deficient (KO; *i.e*., B6.*Slc10a2*^-/-^) mice. Intact extracts were normalized to add 200-300 μM total BAs of each WT or KO extract, based on colorimetric kit analysis, as in (D); identical dilutions of parallel BA-depleted extracts were used. Numbers of differentially expressed genes (red, upregulated; blue, downregulated), for each condition *vs*. vehicle (DMSO), are indicated by color matched text. Data were pooled from 2-independent experiments (1 or 3 biological replicates each) using siLC extracts from separate wild type or Asbt-deficient mice.

**(F)** Mean normalized *Fgf15* mRNA expression [+ SEM; *n* = 3-4; expressed as transcripts per million (TPM)] in cultured ileal explants, determined by RNA-seq as in (E). \**P* < 0.05, \*\**P* < 0.01, one-way ANOVA with Holm-Sidak correction for multiple comparisons or unpaired two-tailed Student’s *t* test.

## Methods

### Mice

C57BL/6J (B6) mice (stock no: 000664) were purchased from the Jackson Laboratory (ME, USA). B6-derived *Slc10a2*^-/-^ (B6.*Slc10a2*^-/-^) mice were obtained from Dr. Paul Dawson (Emory University, GA, USA) ^43^. Most mice used in these experiments were female, to avoid fighting by males of different litters; animals were weaned at 4-5 weeks old, co-housed at 8-10 weeks old and sacrificed at 29-32 weeks old unless otherwise stated. All breeding and experimental use of animals was conducted in accordance with protocols approved by IACUC committees at Scripps Florida or Dartmouth College.

### Bile acid metabolomics

Mice were separated into individual cages and fasted for 4 hr prior to sample collection. Samples (siLC, smvB, feces, pB) were collected, weighed, and stored at -80°C until sent for BA quantification. siLC was harvested from the whole SI. pB from submandibular cheek bleeds with 5mm Goldenrod animal lancet (Medipoint, NY, USA) were collected into MiniCollect 1mL Lithium Heparin coated tubes (Greiner Bio-One, NC, USA). smvB was subsequently collected from surgically open live mice under 3% isoflurane nose cone administered anesthesia using 5ml syringe attached to a 30G mouse PE CSF collection lines (SAI, IL, USA) coated in Lithium Heparin. siLC was harvested from the whole SI following EA-34000 SMARTBOX CO_2_ euthanasia (E-Z System Inc., PA, USA) immediately after smvB collection. BA quantification analyses were performed by Creative Proteomics (NY, USA). Briefly, for blood, 20 μL samples were mixed with 80 μL of the internal standard (IS) solution (50% methanol/50% water containing 0.01% formic acid, deuterated BAs) and 900 μL water. The deuterated BAs were d4-CA, 2,2,4,4-d4-UDCA, d4-CDCA, d4-DCA, d4-LCA, d4-(t)UDCA, d4-(t)CA, 2,2,4,4-d4-(g)UDCA, d4-(g)CA, d4-(t)CDCA, d6-(t)DCA, d4-(g)CDCA, 2,2,4,4-d4-(g)DCA, 2,2,4,4-d4-(g)LCA.

After samples were sonicated on ice for 10 min, the mixture was loaded onto a polymeric reversed-phase SPE cartridge (60 mg/mL). After washing with water, BAs were eluted with 1 mL methanol. The flow-through fraction was collected under a 2-psi positive pressure, dried under a nitrogen gas flow, and reconstituted in 80 mL of 50% acetonitrile. For siLC and feces, samples were homogenized in 20 μL of 70% acetonitrile per 1 mg tissue using a Mixer mill MM 400 (Retsch, Haan, Germany) at 30 Hz for 3 min, sonicated in a water bath for 3 min, then centrifugation. The supernatant was diluted with IS solution containing. A mixture of standards containing all the target BAs was dissolved in an IS solution containing 14 D-labeled BAs and used for calibration. An Agilent 1290 UHPLC system (Agilent, CA, USA) coupled to a Sciex 4000 QTRAP mass spectrometer (Sciex, MA, USA) was used. The MS instrument was operated in the multiple reaction monitoring mode with negative-ion detection. A Waters BEH C_18_ column (2.1*150 mm, 1.7 mm) was used, and the mobile phase was (A) 0.01% formic acid in water and (B) 0.01% formic acid in acetonitrile for binary-solvent gradient elution. For further analyses with similar matrix algorithm and Pearson correlation, Morpheus (https://software.broadinstitute.org/morpheus) was utilized.

### Intestinal explant cultures

The terminal 6 cm of ileum, or 3 cm of the proximal colon, was harvested from 8-18 weeks old male B6 mice after fasting for 4 hr. Fat was removed and tissues were opened longitudinally. Luminal contents were gently flushed with ice-cold PBS, and fragments were first cut into 0.5 cm pieces along the vertical axis; these pieces were subsequently cut into halves along the horizontal axis and placed in ice-cold PBS prior to culture. Fragments containing peyer’s patches were not used. 1 cm x 1 cm square sponges (Surgifoam; Ethicon, NJ, USA) were soaked in 1 ml of serum-free Advanced DMEM/F12 (Gibco, MA, USA) containing 100 U/ml penicillin, 100 μg/ml streptomycin, 10 mM HEPES, 2 mM GlutaMAX (Gibco) in 12-well plate wells. 0.5 cm x 1 cm sponges and 0.5 ml of medium were used for cultures in 24-well plates. Air in sponges was released by applying manual pressure. Intestinal fragments were then dried gently on kimwipes, opened apical side up, and placed on media-soaked sponge. Plates were Incubated at 37°C with 5% CO_2_. In some experiments, 0.2-0.6% dimethyl sulfoxide (DMSO), 50-400 μM (t)CDCA (Sigma-Aldrich, MO, USA), 200 μM GW4064 (MedChemExpress, NJ, USA) and/or 10 μM odevixibat (MedChemExpress) were used.

For experiments where cultured explants were treated with endogenous siLC extracts, siLC was collected from whole SI of 8-to 9-wk old B6 or B6.*Slc10a2*^-/-^ mouse cohoused for at least 4 weeks, weighed and stored at -20°C. Samples were homogenized in 1 ml of a mixture of 75% acetonitrile, 10% methanol, 15% water with 0.5 g of 2.7 mm dia glass beads (BioSpec Products, OK, USA) and SPEX SamplePrep 1600 miniG (Cole-Parmer, NJ, USA) at 1,500 strokes/min for 1 min with 3 sets. After incubated on a nutator (60 min, RT), samples were centrifuged (16,000 g, 10 min, 4°C) and the supernatants were harvested. These steps were repeated using the pellets from the first extraction. The pooled supernatants were dried up by Savant SpeedVac SPD120 (Thermo Fisher Scientific, MA, USA) and redissolved in DMSO. To deplete BAs, the samples were incubated with 19-fold volume of 50 mM cholestyramine (Sigma-Aldrich)-containing PBS for 60 min on a shaker (200 rpm, RT). After centrifugation (16,000 g, 10 min, 4°C), the supernatants were harvested. As cholestyramine-untreated groups, the samples were diluted in a 19-fold volume of PBS. Total bile acids assay (Diazyme Laboratories, CA, USA) and BioTek Epoch Microplate Spectrophotometer (Agilent Technologies, CA, USA) were used to measure total BA concentrations to determine the volume of cholestyramine-untreated extract needed to add for a final concentration of ∼300 μM BAs in the ileal explant culture media. The same volume of cholestyramine-treated extract was used for the BA-depleted media.

### Gene expression assays

*qPCR*: Total RNA was isolated from cultured or *ex vivo* tissues using TRIzol (Invitrogen, MA, USA) or RNeasy Mini Kit (Qiagen, MD, USA; with on-column DNase I treatment) after mechanical disruption with 0.5 g of 2.7 mm dia glass beads and SPEX SamplePrep 1600 miniG (1,500 strokes/min, 1 min, 2 sets). RNA was quantified on a NanoDrop 2000 (Thermo Scientific, MA, USA). qPCR was performed with SuperScript III Platinum One-Step qRT-PCR Kit (Invitrogen) and commercial Taqman primer/probe sets (Applied Biosystems, MA, USA) on a QuantStudio 6 Pro (Applied Biosystems). Probes included: *Actb* (Mm00607939_s1) and *Fgf15* (Mm00433278_m1). Data were analyzed with Design & Analysis (Applied Biosystems).

#### Nanostring

Tissues (spleen, liver, colon, terminal ileum) were harvested post-mortem from mice after a 4 hr fasting period and blood (peripheral, portal) collection, as above. Total RNA was isolated using RNeasy Mini kits with on-column DNase I treatment. Gene expression was analyzed using a custom codeset run on an nCounter Pro system (Nanostring, WA, USA) as per manufacturer’s instructions. Data were analyzed with nSolver software (Nanostring).

### Bioinformatics

#### Metagenomics

feces were collected from individually-housed mice after a 4 hr fasting time and stored at -80°C prior to shipment to TransnetYX (TN, USA) for library preparation and sequencing. DNA was extracted using Qiagen DNeasy 96 PowerSoil Pro QIAcube HT kits (Qiagen). Sequencing libraries were prepared with Watchmaker DNA Library Prep kits with Fragmentation (Watchmaker, CO, USA) and sequenced using the Illumina NovaSeq instrument (Illumina, CA, USA) with the shotgun sequencing method (a depth of 2 million 2x150 bp read pairs). Acquired data were analyzed with the One Codex database. Metagenomics reads were processed using atlas v.2.13.0^84^. In short, using tools from the STAR suite v.2.7.2b^85^, to filter out contaminations from the mouse host genome. Reads were error corrected and merged before assembly with metaSpades v.3.15.3^86^. Contigs were binned using maxbin2 v.2.2^87^. The predicted MAGs were clustered (95% average nucleotide identity) resulting in 34 representative genomes (referred later as genomes). Genes of each genome were predicted using eggNOG v.2.1^87^. α- and β-diversity of metagenome communities were calculated using the relative abundance of MAGs from the atlas workflow with vegan R package v.2.6-4^88^.

*RNA-seq*: bulk RNA-seq analysis was performed on wild type (B6) ileal fragments cultured with DMSO, (t)CDCA, or endogenous BA extracts with or without CME treatment from B6 or B6.*Slc10a2*^-/-^ mice, as described above. Total RNA was isolated with TRIzol (Invitrogen, MA, USA) after mechanical disruption with 0.5 g of 2.7 mm dia glass beads and SPEX SamplePrep 1600 miniG (1,500 strokes/min, 1 min, 2 sets), and quantified using a Qubit Fluorometer (Invitrogen). RNA quality was evaluated using a 5200 Fragment Analyzer System (Agilent Technologies). QuantSeq 3’ mRNA-Seq V2 Library Prep Kit with UDI (Lexogen, Vienna, Austria) was used for cDNA library preparation from 100 ng total RNA. Libraries were pooled and sequenced using 10 M, single-end, 100-bp forward strand reads per sample on NextSeq2000 (Illumina, CA, USA). DNA alignment, quantification, and annotation were performed using the Dartmouth Analytics Core RNAseq pipeline. Quality control was evaluated using fastQC. Reads trimmed using Cutadapt were aligned to the NCBI mm39 mouse reference genome using the HSIAT2 aligner. Sequencing metrics and ribosomal RNA identification were performed using the Picard Toolkit, and a report was created using MultiQC. Samples with low sequencing depths were removed. Differential gene expression (DGE) analysis was performed with DESeq2 (*P*adj < 0.05) to enumerate genes differentially expressed in each condition, relative to DMSO. Gene symbols were annotated by Entrez gene ID using the org.Mm.eg.db database. ggplot2 was used to visualize data.

### Mathematical modeling of BA circulation and metabolism

BA flux through the intestine was modeled as a series of connected compartments^59^. Additional compartments such as liver and gallbladder were not modeled to focus on the effects of reabsorption on the intestinal BA pool. Only 1° and 2° BA species were used as a measure of the BA pool diversification. BA concentrations were governed by differential equations accounting for synthesis, transit, reabsorption, and conversion reactions. Units were provided in molar amounts, and the fluxes between the compartments were balanced. Reaction rates that fit our experimental data qualitatively were numerically calculated through parameter optimization. The compartment volumes were required to quantitatively compare with the actual rates determined in the experiment to convert BA units to concentrations. Changes in BA x amount in compartment I were given by:

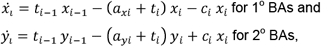

where *t*_*i*-1_ was the input rate of BAs transiting from the previous compartment; was the removal rate of BAs from the compartment; *a*_*xi*_ and *a*_*yi*_ were the **reabsorption** rates of 1°- and 2° BAs, respectively; c represented the metabolization rate of 1° BAs into 2° BAs. In the first compartment, the rate *s* of 1° BA synthesis plus the sum of all reabsorbed BAs were used as the input, instead of BAs input from recirculation:

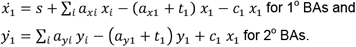

The whole intestine was divided into 15 compartments, (10 in SI, 5 in colon). BA intestinal transit speed was set to rates *t*_*si*_ in SI compartments 1-8, reduced to *t*_*il*_ in SI compartments 9-10, and decreased further to *t*_*co*_ in colon compartments 11-15. The colonic transiting speed was considered 10-fold slower than that in the SI, based on the transit of technetium-labeled activated charcoal DTPA in mice^89^. Rates of passive absorption were considered different in the SI and colon, with rates *p*_*si*_ and *p*_*co*_, respectively. Active ileal reabsorption was considered in simulations of wild type B6 mice, *i.e*., *a*_8,9_ = *p*_*si*_ +*a*_*act*_. The reabsorption rates of 1°- and 2° BAs were considered same in the initial simulation. 1° → 2° BA metabolism was considered only in the colon, *i.e*.,

MatLab solver *ode113* was used to implement simulations of this system, and an optimization of absorption and conversion rates were performed to qualitatively fit our experimental data. For the optimization, a single set of parameters [*a*_*act*_ *p*_*si*_ *p*_*co*_ *c*_*co*_] was considered and the system was simulated twice in the presence absence of active reabsorption. fminsearch and the sum of squared errors were used as a cost function. Code is available at github.com/schultz-lab/Bile_Acids.

Since elimination of active reabsorption alone was not sufficient to explain the diversification of the BA pool in B6.*Slc10a2*^-/-^ mice, an increase of bacterial BA metabolism was implemented as a possible mechanism to increase the amount of 2° BA in the absence of active ileal absorption. Different rates of bacterial BA metabolization were used, *c*_*wt*_ and *c*_*ko*_for B6 and B6.*Slc10a2*^-/-^ mice in our optimization.

Experimental data used in optimization:

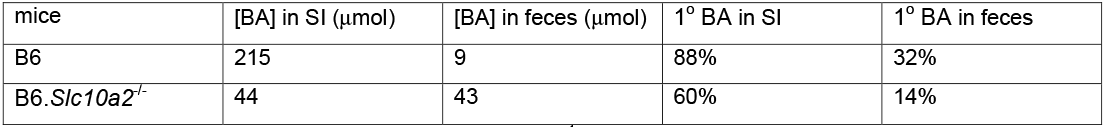

Parameters used in simulations (rates given in min^-1^):

| $t_{si}$ | $t_{il}$ | $t_{co}$ | $a_{act}$ | $p_{si}$ | $p_{co}$ | $c_{co}$ | $c_{wt}$ | $c_{ko}$ | $K_x$ | $K_y$ |
| --- | --- | --- | --- | --- | --- | --- | --- | --- | --- | --- |
| 0.1 | 0.05 | 0.01 | 0.29 | 0.025 | 0.0012 | 0.0030 | 0.0010 | 0.0090 | 1.20 | 4.29 |

### Anaerobic bacterial culture

Cryopreserved mouse fecal pellets were thawed, weighed and resuspended in sterile PBS supplemented with 10 mM L-cysteine in a ratio of 1 to 10 w/v. Following homogenization, the samples were inoculated into MiPro medium^90^ in a sterile 12-well plate to a final v/v ratio of 2%. Each culture was passaged anaerobically for 24 hours at 37°C in a Whitley A55 anaerobic chamber (Don Whitley Scientific, Victoria 412 Works, UK) with gas composition of 10% CO_2_, 10% H_2_, 80% N_2_. The culture was analyzed as follows: (*i*) CFU/mL were calculated by plating aliquot of the planktonic culture on sheep blood agar that was then incubated at 0% or 21% oxygen, (*ii*) the cell pellets were collected and stored in RNAprotect (Qiagen, MD, USA) at -80°C for DNA extraction, 16S rRNA gene amplicon library sequencing and metagenomic sequencing, and (*iii*) culture supernatants were filter-sterilized and stored at -80°C for subsequent analysis of bile acid metabolites.

### Statistical analysis

Statistical analyses were performed with Prism (GraphPad Software, MA, USA). *P* values were determined as shown in the figure legends. Data are expressed as the mean + SEM.

